# Neural correlates of retrieval success and precision: an fMRI study

**DOI:** 10.1101/2024.06.10.598309

**Authors:** Mingzhu Hou, Paul F. Hill, Ayse N. Z. Aktas, Arne D. Ekstrom, Michael D. Rugg

## Abstract

Prior studies examining the neural mechanisms underlying retrieval success and precision have yielded inconsistent results. Here, their neural correlates were examined using a memory task that assessed precision for spatial location. A sample of healthy young adults underwent fMRI scanning during a single study-test cycle. At study, participants viewed a series of object images, each placed at a randomly selected location on an imaginary circle. At test, studied images were intermixed with new images and presented to the participants. The requirement was to move a cursor to the location of the studied image, guessing if necessary. Participants then signaled whether the presented image as having been studied. Memory precision was quantified as the angle between the studied location and the location selected by the participant. A precision effect was evident in the left angular gyrus, where BOLD activity covaried across trials with location accuracy. Multi-voxel pattern analysis also revealed a significant item-level reinstatement effect for high-precision trials. There was no evidence of a retrieval success effect in the angular gyrus. BOLD activity in the hippocampus was insensitive to both success and precision. These findings are partially consistent with prior evidence that success and precision are dissociable features of memory retrieval.

## 1. Introduction

Episodic memory refers to conscious memory of personally experienced, unique events, typically defined by their spatiotemporal context (Tulving, 1983). Successful episodic memory retrieval (recollection) is typically defined as the retrieval of qualitative (usually contextual) information about a prior event. A different memory signal, typically referred to as familiarity, is held to support a sense of past occurrence without providing access to qualitative information about the event.

Most studies examining recollection have employed categorical memory measures to assess whether contextual information associated with a studied item can be remembered. For example, in studies employing the ‘remember/know’ paradigm (e.g. Duarte et al., 2010; Wheeler and Buckner, 2004), recollected items are defined as those that are both correctly recognized and accompanied by a declarative memory for some aspect of the prior event. In studies using source memory tests, recollection is indexed by successful retrieval of diagnostic contextual information belonging to the studied event, such as the studied color or location of a test item (e.g. Diana et al., 2010; Slotnick et al., 2003). In studies employing associative memory procedures (e.g. de Chastelaine et al., 2016; Giovanello et al., 2004), recollection is operationalized by successful recall of the relational information linking two study items.

Although widely employed, studies using categorical memory measures provide limited information about memory *precision* – the degree of correspondence between the originally experienced and the remembered event. In some recent studies the constructs of retrieval success (the accessibility of an encoded event) and precision have been separately estimated by the use of what Harlow and Donaldson (2014) termed a *positional response accuracy* paradigm (e.g. Harlow and Donaldson, 2014; Harlow and Yonelinas, 2016; Korkki et al., 2023; Richter et al., 2016). In these studies, memory performance was quantified by a continuous metric that reflected the difference between a specific feature of a study event and the corresponding recollected feature (e.g. the difference between the location of a studied item and its recalled location). Trial-wise distance metrics are fit by a two-component mixture model comprising a rectangular distribution that models the probability of a random guess (i.e. unsuccessful retrieval), and a circular Gaussian distribution that models the probability of successful retrieval with varying precision.

The proposal that retrieval success and precision can be dissociated has received support from several behavioral studies. For example, Harlow and Yonelines (2016) reported that participants can subjectively differentiate success from precision on a trial-by-trial basis. Additionally, retrieval practice has been reported to increase the likelihood of retrieval success but not the precision of the retrieved memories (Sutterer and Awh, 2016). In other studies, it was reported that the likelihood of retrieval success declines faster than precision as a function of retention interval (Berens et al., 2020). By contrast, precision is more sensitive than success to healthy aging (Gellersen et al., 2024; Korkki et al. 2020; Nilakantan et al., 2018) and damage to the hippocampus (Nilakantan et al., 2018).

To our knowledge, only three fMRI studies have employed the positional response accuracy procedure to separately examine the neural correlates of retrieval success and precision (Cooper et al., 2017; Korkki et al., 2023; Richter et al., 2016; for evidence from functional connectivity, see Cooper and Ritchey, 2019). The studies each employed a region-of-interest (ROI) approach to examine neural activity in the left angular gyrus (AG) and hippocampus, and yielded inconsistent findings. Richter et al. (2016) reported that, in a sample of healthy young adults, retrieval success was exclusively associated with enhanced activity in the hippocampus, while AG activity covaried on a trial-wise basis with precision. This general pattern of findings was replicated in participants with autism spectrum disorder and neurotypical controls in Cooper et al. (2017). However, Korkki et al. (2023) reported that fMRI BOLD activity in the AG and hippocampus was sensitive to both success and precision. The finding for the hippocampus is consistent with the results of the lesion deficit study of Nilakantan et al. (2018), in which it was reported that patients with medial temporal lobe lesions that included the hippocampus demonstrated lower precision estimates than did patients with exclusively extra-hippocampal lesions (see also Borders et al., 2022).

In light of the inconsistent findings from these prior fMRI studies, here we sought to further examine the neural correlates of retrieval success and precision. In addition to examining success- and precision-related neural activity in hippocampal and AG, we employed an exploratory whole brain analysis. Unlike prior studies that assessed memory precision for multiple features (Cooper et al., 2017; Korkki et al., 2023; Richter et al., 2016), we employed a study design focused specifically on precision for spatial location, mirroring the approach taken by Harlow and colleagues (Harlow and Donaldson 2014; Harlow and Yonelinas, 2016), and Nilakantan et al. (2018). In a significant extension of prior studies, we employed both univariate and multi-voxel pattern similarity analysis (PSA) to examine the neural correlates of memory precision. The PSA was predicated on prior reports that item-wise retrieval-related ‘reinstatement’ (the overlap between across-voxel patterns of BOLD activity elicited at study and test) is especially prominent in the AG (Kuhl and Chun, 2014), where it has been reported to covary with a metric of memory specificity (Lee et al., 2019). Multi-voxel reinstatement effects have also been reported in the hippocampus, where they covary with retrieval accuracy (e.g. Liang and Preston, 2017; Tompary et al., 2016). Accordingly, in the present study we contrasted metrics of item-level reinstatement derived from the AG and hippocampus according to the precision of the associated memory judgments, with the prediction that reinstatement should be stronger for high-than for low-precision judgments.

## 2. Methods

### 2.1 Participants

The study sample comprised cognitively healthy young adults (N = 23, mean age = 24 yrs, age range = 18-30 yrs, 12 female). The sample size was selected to be comparable with the samples employed in previously reported fMRI studies of memory precision in young adults (Richter et al., 2016, N = 21; Korkki et al., 2023, N = 23). Each participant was right-handed, had normal or corrected-to-normal vision, no history of neurological or psychiatric illness, and was not taking prescription medications that affected the central nervous system. Informed consent was obtained in accordance with UT Dallas Institutional Review Board guidelines. Participants were compensated at the rate of $30 an hour. Data from four additional participants were excluded due to excessive head movement during the scan (N = 1) and poor memory performance on item recognition (below-chance level of item hit rate, N = 3).

### 2.2 Experimental items

Items comprised 136 images of everyday objects (see Figure 1). Of these images, 102 were employed as study items and an additional 34 images were employed as new items at test.

**Figure 1.**
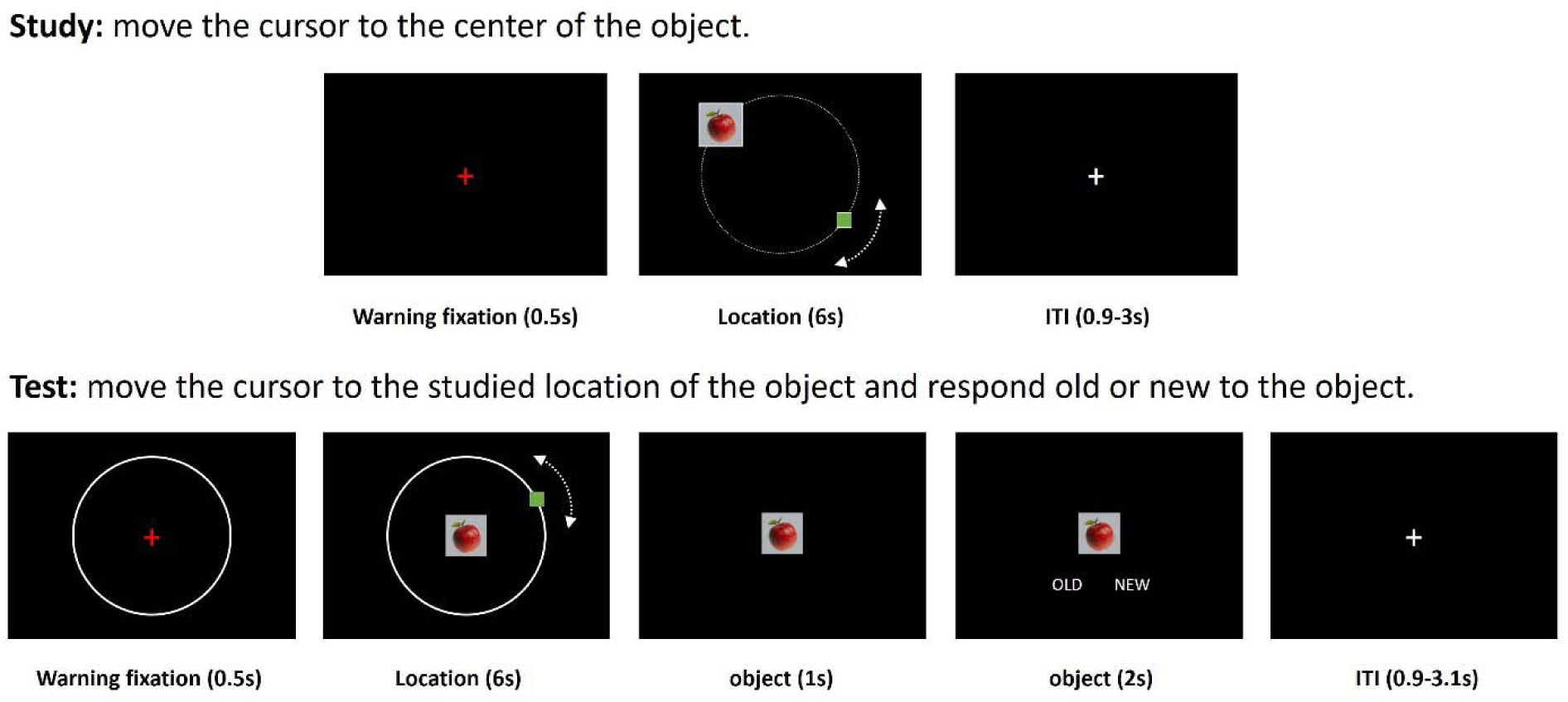
Schematic depiction of the study and test tasks. ITI: inter-trial interval.

### 2.3 Procedure

Figure 1 provides a schematic depiction of the study and test tasks. Study and test items were viewed via a mirror mounted on the scanner head coil that reflected a back projected image (viewing distance 102 cm). There was a single study block followed by two consecutive test blocks. At study, participants viewed 102 object images superimposed on a grey background (visual angle: 1.4 x 1.4 deg). Each image was presented for 6 s at a randomly selected location on a virtual circle (radius = 4.9 deg) for 6 s. The task was to move a small green square (the cursor) along the circle by pressing one of two buttons until it overlapped the image (see Figure 1). Participants were instructed to remember both the image and its location.

Test items comprised a randomly ordered sequence of the studied images intermixed with 34 unstudied images. The images were presented for 6 s at the center of a delineated circle with the same radius as the virtual circle employed during the study phase. The task was to move a cursor, which was randomly located on the circle, to the location that had been occupied by the image at study, guessing if necessary. Participants were instructed to move the cursor to a random location if they did not recognize the image as having been studied. After moving the cursor, they signaled whether the presented object had been studied or unstudied. The ordering of the test images was pseudo-randomized so that no more than 3 old or new images occurred in succession.

Participants responded via a button box. They pressed the buttons under their right index and middle fingers both to move the cursor along the circle and, at test, to respond ‘old’ or ‘new’ to the object image. The mapping between the button presses and the direction of the cursor movement, as well as the old/new judgments, were counterbalanced across participants.

During both study and test, a 30-s break occurred in the middle of the presentation blocks, when a ‘rest’ cue was presented at the center of the screen. Participants were instructed to relax until the cue disappeared, at which point the task continued. Before the study block, and after the 2^nd^ test block, participants also performed a ‘baseline’ task. The procedure was the same as that for the study task except the instructions were to move the cursor to the center of a randomly located ‘X’ character. Data from the baseline task are not reported here.

### 2.4 MRI acquisition and preprocessing

MRI data were acquired with a Siemens PRISMA 3T MR scanner equipped with a 32-channel head coil. Functional scans were acquired with a T2^D^-weighted echoplanar (EP) sequence [TR 1560 ms, TE 30 ms, flip angle 70°, field-of-view (FOV) 220 mm, multiband factor = 2, 50 slices, voxel size 2.5 x 2.5 x 2.5 mm, 0.5 mm inter-slice gap, anterior-to-posterior phase encoding direction]. A T1-weighted image was acquired with an MP-RAGE pulse sequence (TR = 2300 ms, TE = 2.41 ms, FOV 256 mm, voxel size = 1 x 1 x 1 mm, 160 slices, sagittal acquisition). A T2-weighted image was also acquired, focusing on the medial temporal lobe (TR: 3500 ms, TE: 36 ms, FOV: 215 mm, flip angle 120°, voxel size = 0.42 x 0.42 x 0.42 mm, 40 slices). A field map was acquired after the functional scans using a double-echo gradient echo sequence (TE 1/2 = 4.92 ms/7.38 ms, TR = 520 ms, flip angle = 60°, FOV 220 mm, 50 slices, 2.5mm slice thickness). Functional images were acquired during the baseline, study and test tasks, but here we only report findings from the study and test phases.

Data preprocessing was performed with SPM12 implemented in MATLAB R2018b. The functional volumes were field-map corrected, realigned, reorientated to the anterior commissure – posterior commissure line and spatially normalized to the MNI EPI template. The normalized images were resampled to 2.5 mm isotropic voxels and smoothed with a 6 mm Gaussian kernel. Before being entered into univariate subject-wise General Linear Models (GLMs), the functional data from the two test blocks were concatenated using the spm_concatenate.m function. Anatomical images were normalized to an MNI T1 template.

### 2.5 Statistical analyses

#### 2.5.1 Behavioral analyses

For each participant, we computed the proportion of correctly recognized old items (item hit rate) and the proportion of correctly rejected new items (correct rejection rate). Item memory performance (Pr) was computed as p(hit) – p(false alarm).

Across-participants distance error (the difference between the estimated and actual studied location in deg) for item hit trials were fit to a group-level two-component mixture model, using standard mixture modelling as implemented in the MemToolbox (Suchow et al., 2013). The model estimated a rectangular distribution reflecting the proportion of guess responses (g), and a von Mises distribution reflecting accurate location memory with varying precision [indicated by the standard deviation (sd) of the distribution]. Item hit trials were divided into successful and guess trials depending on whether the associated distance errors had a < 5% chance of fitting the group-wise von Mises distribution. To anticipate the results, the cutoff for the location hit vs guess trials in the present study was +/- 47 deg (i.e. location hit trials were those item hit trials with an absolute distance error < 47 deg).

#### 2.5.2 fMRI analyses

##### 2.5.2.1 Whole brain analyses

In the whole brain analyses, we investigated retrieval success effects by contrasting location hit and guess trials. To identify precison effects we examined the relationship between BOLD activity and the trial-wise distance error of location hits. Each of these analyses was based on the same first level participant-wise GLMs, which were taken forward to two separate across-participant GLMs.

For the participant-wise GLMs, a boxcar function spanning the duration of the retrieval cue (6s) was used to model neural activity synchronized to item onset. Three event types of interest were included: location hits (item hit trials with absolute distance error < 47 deg), guesses (item hit trials with absolute distance error > 46 deg) and correct rejections (CRs). For the location hits, a trial-specific measure of memory precision was included as a parametric modulator to estimate the degree to which neural activity varied with precision. Precision was estimated as the absolute distance error in degrees between the judged and the studied location (errors were subtracted from 180 before entering analysis so that higher values indicated higher precision). The series of precision estimates across trials was mean-centered within participants. Mean trial numbers for location hits, guesses and CRs were, respectively, 42 (range 22-88), 30 (8-49) and 28 (20-34). Other events, including rest breaks, false alarms, misses and trials with absent or multiple old/new responses were modelled as events of no interest. The GLMs also included as covariates six regressors modeling motion-related variance (three for rigid-body translation and three for rotation) and 2 constants for means across test blocks. Data from volumes with a transient displacement (relative to the prior volume) of > 1 mm or > 1◦ in any direction were modeled as covariates of no interest.

Parameter estimates derived from the first level GLMs were taken forward to two second-level, group-wise GLMs. In the first model, parameter estimates for the parametric regressor, which reflected the strength of the relationship between neural activity and memory precision for location hits, were subject to a one-sample t-test. In the second GLM, group-wise effects were examined with a repeated measures one-way ANOVA model that employed the factor of trial-type (3 levels: location hits, guesses and correct rejections). Results from both the t-test and the ANOVA were deemed significant if they survived a height threshold of p < 0.001 and a cluster extent threshold FWE corrected to p < 0.05.

For each significant cluster identified by the 3-level ANOVA model, parameter estimates for location hits, guesses and CRs were extracted from voxels within a 5mm radius of the peak of each cluster and averaged. These averaged estimates were entered into a trial type (location hits, guesses, correct rejections) x region ANOVA. Note that we do not report the main effect of trial type in this analysis since it is a foregone conclusion.

##### 2.5.2.2 Region of interest (ROI) analyses

Given our *a priori* interest in the left AG and hippocampus (see Introduction), we conducted ROI analyses in these two regions. For the left AG, we selected the peaks reported in two previous studies to show AG precision effects (MNI coordinates: -54, -66, 33 in Korkki et al., 2023, and -54, -54, 33 in Richter et al., 2016). We did not consider the findings from Cooper et al. (2017) because the effects in that study were identified across autistic and neurotypical participants. Parameter estimates were extracted and averaged within a 5 mm radius of each AG peak. Given the prior findings suggesting that precision- and memory-related hippocampal effects were maximal in the vicinity of the anterior hippocampus (Korkki et al., 2023; Ritcher et al., 2016), we employed the anatomically defined anterior hippocampus as an ROI. Parameter estimates were extracted from left and right anterior hippocampal masks (hippocampus anterior to y =-22 in MNI space, Poppenk et al., 2013) that had been manually traced on the group mean T1 image (we did not employ the hippocampal peaks reported in the above-mentioned studies because two of these peaks were located outside of the hippocampus as it was identified in our mean T1 image. However, the findings did not differ in secondary analyses that employed these peaks as the centers of 3 mm radius spherical ROIs). For both the AG and hippocampus, precision effects were evaluated with one-sample t-tests conducted on the beta estimates associated with the parametric regressor. Retrieval success effects were examined by the contrast between the beta estimates associated with location hit and guess trials.

##### 2.5.2.3 Additional ROI analyses

To complement the findings from the parametric modulation analyses, we conducted additional analyses that examined precision effects categorically by contrasting the BOLD activity elicited on high- vs. low-precision hit trials. We constructed subject-wise GLMs in which three event types were modeled: high-precision hits (item hit trials with absolute distance error < 16 deg), low-precision hits (item hit trials with absolute distance error ranging from 25-40 deg) and guesses (item hit trials with absolute distance error > 59 deg). Thus, we employed an equivalent accuracy range (15 deg) for the high and low-precision trials. A gap was included between the boundary of these two trial types, and between the low-precision trials and the model-derived cutoff (47 deg) to accentuate any categorical differences. In the case of the low precision trials, we adopted a lower cut-off of 40 deg rather than 46 deg to minimize the likelihood of including trials that more properly belonged to the guess distribution. For the same reason, guess trials were defined as those where distance error was substantially below the estimated 47 deg cut-off. All other trial types were modeled as events of no interest. Parameter estimates for the different categories were extracted from the AG and hippocampal ROIs and subject to pairwise t-tests. The precision effect was operationalized by the contrast between high vs. low precision hit trials. Retrieval success effects were assessed by contrasting high and low-precision location hits with guess trials (contrast weights of +0.5, +0.5, -1).

##### 2.5.2.4 Multi-voxel pattern similarity analyses

In addition to the univariate analyses described above, we conducted PSA to estimate item-level reinstatement for location hits. This was accomplished by examining the correspondence between patterns of neural activity elicited during the encoding and test phases of the experiment. As noted in the Introduction, if precision is dependent on the strength of the reinstatement of encoding-related neural activity, location hits associated with high precision should exhibit a greater trial-wise reinstatement effect than low-precision trials, indicative of retrieval of higher fidelity event-specific information.

For each participant, single-trial beta estimates from the study and test phases were derived from first level GLMs that implemented the least squares all (LSA) approach to single trial estimation (Abdulrahman and Henson, 2016; Mumford et al., 2012). Neural activity elicited by the study items was modeled as a 6 s duration boxcar. Each study item was modeled as a separate event of interest, while the 30 s rest period that occurred midway through the study block, the six motion regressors, and a constant modeling the mean BOLD signal in the block were included as covariates of no interest. An analogous single-trial GLM was employed for the test blocks.

PSA was conducted on three ROIs: the left and right anterior hippocampus (note that equivalent results were obtained from an ROI encompassing the entire hippocampus) and an anatomically defined ROI that encompassed the anterior (PGa) and posterior (PGp) subregions of the AG, as defined by the anatomy toolbox v3.0 (Eickhoff et al., 2005, 2006, 2007). To examine reinstatement for test trials associated with different levels of precision, we segregated the trials according to the three memory judgments described in section 2.5.2.3, namely, high-precision location hits, low-precision location hits and guesses. For each trial belonging to each class of memory judgment, encoding-related trial-wise parameter estimates were extracted from each voxel falling within a given ROI. Retrieval-related single-trial parameter estimates were extracted in a similar manner.

Item-level reinstatement for a given trial was operationalized as the difference between within-item and across-item study-test similarity (cf. Hill et al., 2021). Within-item similarity was calculated as the across-voxel Fisher-z transformed correlation between a given study trial and its corresponding test trial. Across-item similarity was computed as the mean of the Fisher-z transformed correlations between the same study trial and all other test trials belonging to the same class of memory judgment. For each participant, within-between similarity estimates were averaged across all of the trials belonging to each class of judgment, providing three summary measures per participant. One sample t-tests were employed to assess whether, across participants, these measures differed reliably from zero. To examine our prediction concerning the association between reinstatement strength and memory precision (see Introduction), planned comparisons were employed to assess whether reinstatment effects differed between high and low precision trials.

Statistical analyses were conducted with SPSS 27.0. Non-sphericity between the levels of repeated measures factors in the ANOVAs was corrected with the Greenhouse-Geisser procedure (Greenhouse and Geisser, 1959). Significance levels for all tests were set at p < 0.05.

## 3. Results

### 3.1 Behavior

Across participants, mean item memory performance (Pr) was 0.57, sd = 0.20, with a mean item hit rate of 0.76, sd = 0.11 and a correct rejection rate of 0.81, sd = 0.13.

Figure 2 shows the distribution of retrieval distance errors for item hit trials pooled across participants (N = 1777 trials). The two-component mixture model yielded an estimated proportion of guess responses (g) of 0.49, and a standard deviation for the von Mises distribution of 23.6 deg (equivalent to a Kappa concentration parameter of 6.43). The cut-off point, which reflected the > 0.05 probability that an item hit trial fell inside the von Mises distribution was +/- 47 deg as estimated using the ‘vonmisescdf’ function from https://www.paulbays.com/toolbox/. We also applied the mixture model on a participant-wise basis. Across participants, the mean guess rate was 0.43, sd = 0.20. The mean standard deviation for von Mises distribution was 28.88, sd = 14.02 (Kappa of 5.38, sd = 2.59).

**Figure 2.**
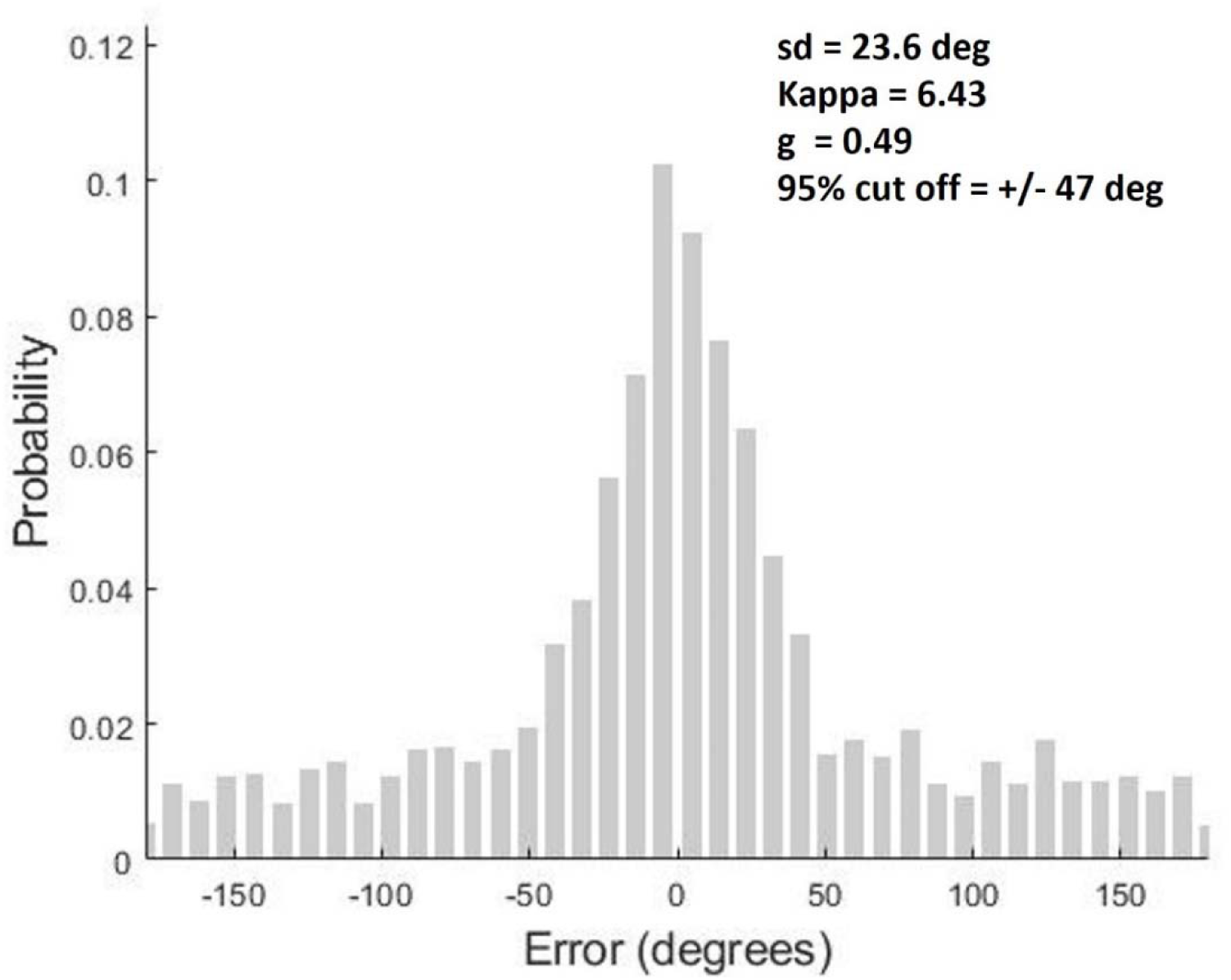
Distribution of distance errors for item hit trials across participants (N = 1777).

### 3.2 fMRI results

#### 3.2.1 Whole brain analyses

At our pre-defined thresholds (see Methods) the parametric modulation analysis did not identify any cluster that demonstrated a precision effect. By contrast, the one-way ANOVA model that employed the factor of trial type (location hits, guesses and correct rejections) revealed a significant main effect in multiple regions, including bilateral dorsal parietal cortex (DPC) extending into the precuneus and posterior cingulate, premotor cortex, left dorsal prefrontal cortex, and left and right AG (see Table 1 and Figure 3). The left AG cluster was closely adjacent to one of the *apriori* defined AG ROIs (MNI coordinates: -54, -66, 33, as reported by Korkki et al., 2023).

**Figure 3.**
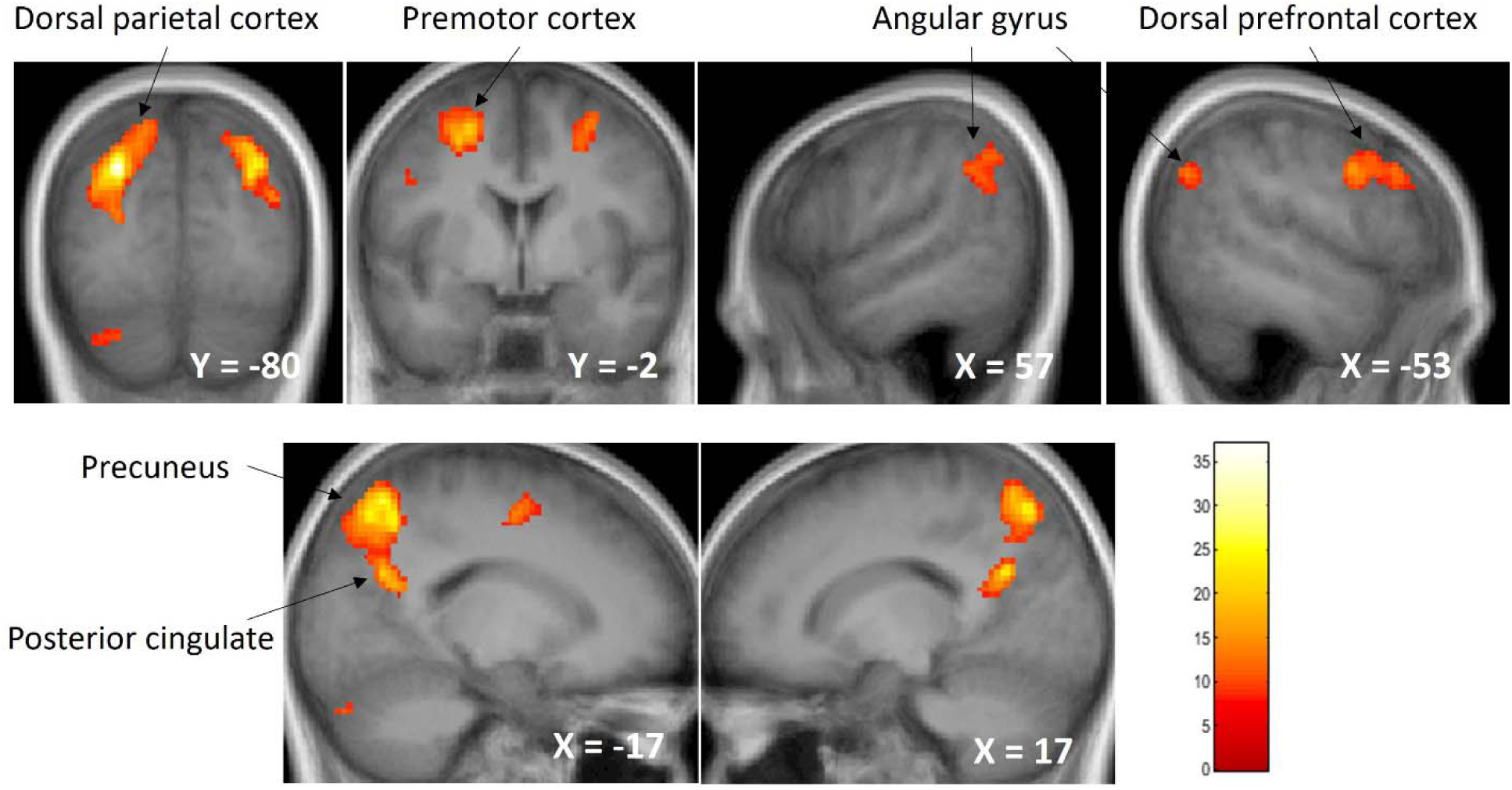
Clusters demonstrating trial type effects in the whole brain analyses.

**Table 1.** Regions demonstrating significant trial type effects in the whole brain analysis. The MNI coordinate of the peak of each cluster is listed.

| Region | MNI |  |  | k | z |
| --- | --- | --- | --- | --- | --- |
|  | x | y | z |  |  |
| L dorsal parietal cortex | -26 | -62 | 52 | 2356 | 6.15 |
| R dorsal parietal cortex | 14 | -72 | 52 | 1223 | 5.45 |
| L premotor cortex | -23 | -2 | 52 | 357 | 4.78 |
| R premotor cortex | 30 | 3 | 55 | 199 | 4.48 |
| L dorsal prefrontal cortex | -56 | 26 | 32 | 329 | 4.45 |
| L AG | -56 | -70 | 32 | 61 | 4.31 |
| R AG | 62 | -52 | 38 | 170 | 4.07 |
| L cerebellum | -20 | -84 | -38 | 61 | 3.88 |

Parameter estimates were extracted from the cortical clusters listed in Table 1 (see Figure 4) and subjected to a trial type (3) x region (7) ANOVA. The ANOVA gave rise to a main effect of region (F_2.78, 61.16_ = 36.90, p < 0.001, partial η^2^ = 0.63), and a trial type x region interaction (F_5.21, 114.64_ = 30.64, p < 0.001, partial η^2^ = 0.58), indicating that the pattern of effects differed across regions. Pairwise t-tests were conducted among the different trial types for each region. Results of the t-tests are summarized in Table 2. As is evident from the table, bilateral DPC and right premotor cortex demonstrated reliable retrieval success (hit > guess) effects. In all other regions except for the left and right AG, the BOLD activity elicited on location hit and guess trials was greater than that elicited by correct rejections. These effects were reversed in left and right AG.

**Figure 4.**
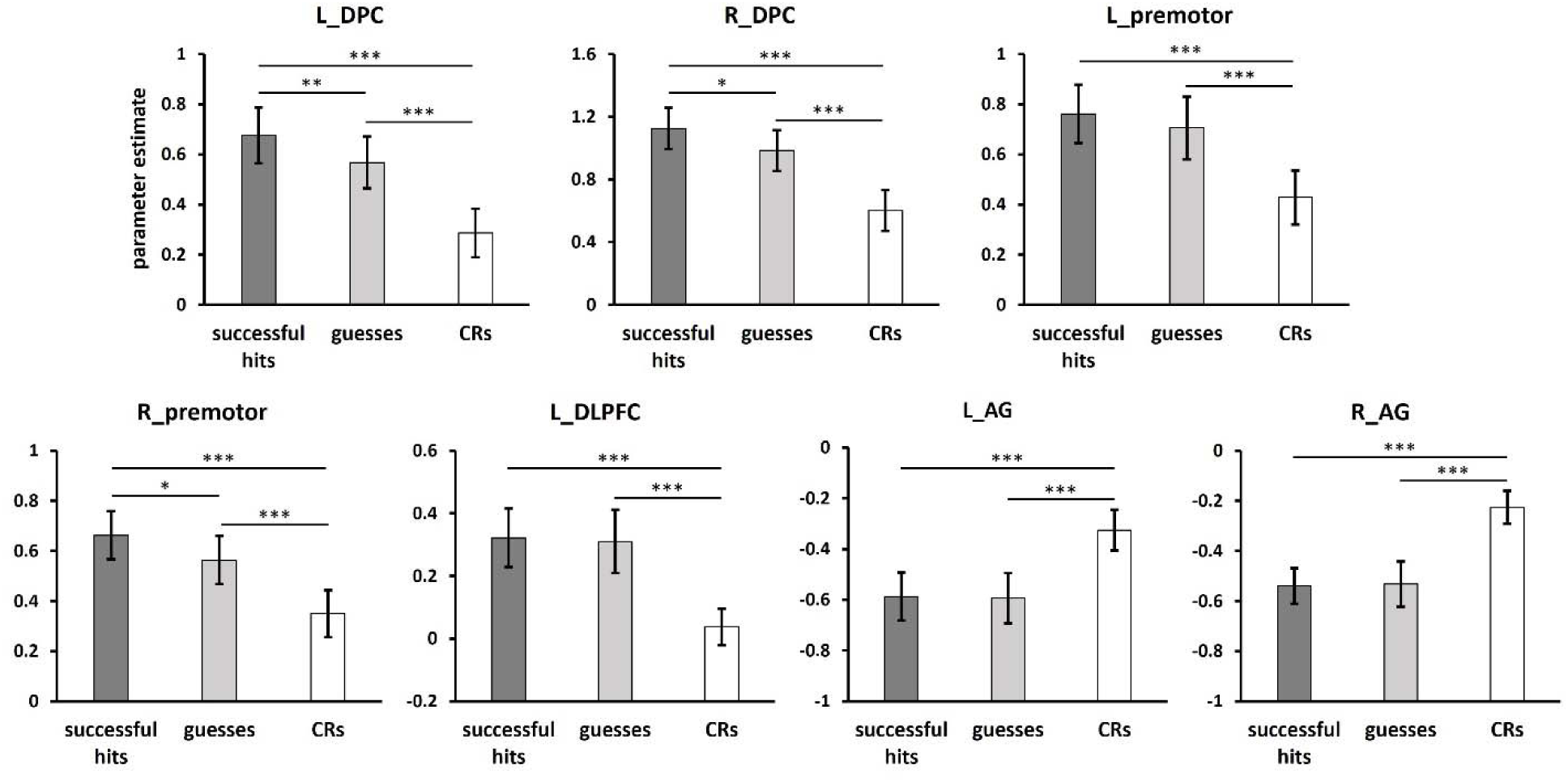
Parameter estimates extracted from the cortical regions identified by the whole brain analysis. Error bars indicate standard errors of the mean. DPC: dorsal partial cortex; premotor: premotor cortex; DLPFC: dorsolateral prefrontal cortex; AG: angular gyrus. *p < 0.05; **p < 0.01; ***p < 0.001.

**Table 2.** P values of pairwise t-tests among different trial types from regions revealed by whole-brain analyses.

|  | L_DPC | R_DPC | L_premotor | R_premotor | L_DLPFC | L_AG | R_AG |
| --- | --- | --- | --- | --- | --- | --- | --- |
| Location hit vs. guess | **0.005** | **0.045** | 0.094 | **0.029** | 0.76 | 0.89 | 0.87 |
| Location hit vs. CR | **< 0.001** | **< 0.001** | **< 0.001** | **< 0.001** | **< 0.001** | **< 0.001<sup>†</sup>** | **< 0.001<sup>†</sup>** |
| Guess vs. CR | **< 0.001** | **< 0.001** | **< 0.001** | **< 0.001** | **< 0.001** | **< 0.001<sup>†</sup>** | **< 0.001<sup>†</sup>** |

Note. CR: correct rejection trials. DPC: dorsal parietal cortex; premotor: premotor cortex;
DLPFC: dorsolateral prefrontal cortex; AG: angular gyrus. <sup>†</sup> CR > location hit and guess.

#### 3.2.2 ROI analyses

As described in section 2.5.2.1, effects of precision and retrieval success were examined in specific AG and hippocampal ROIs. Turning first to the AG, we identified a significant precision effect in the AG region reported by Korkki et al. (2023) (t_22_ = 2.45, p = 0.023, Cohen’s d = 0.51), while the effect in the ROI reported by Richter et al. (2016) failed to attain significance (t_22_ = 1.99, p = 0.059, Cohen’s d = 0.42). No precision effects could be identified in either the left or right hippocampus (ps > 0.52, see Figure 5).

**Figure 5.**
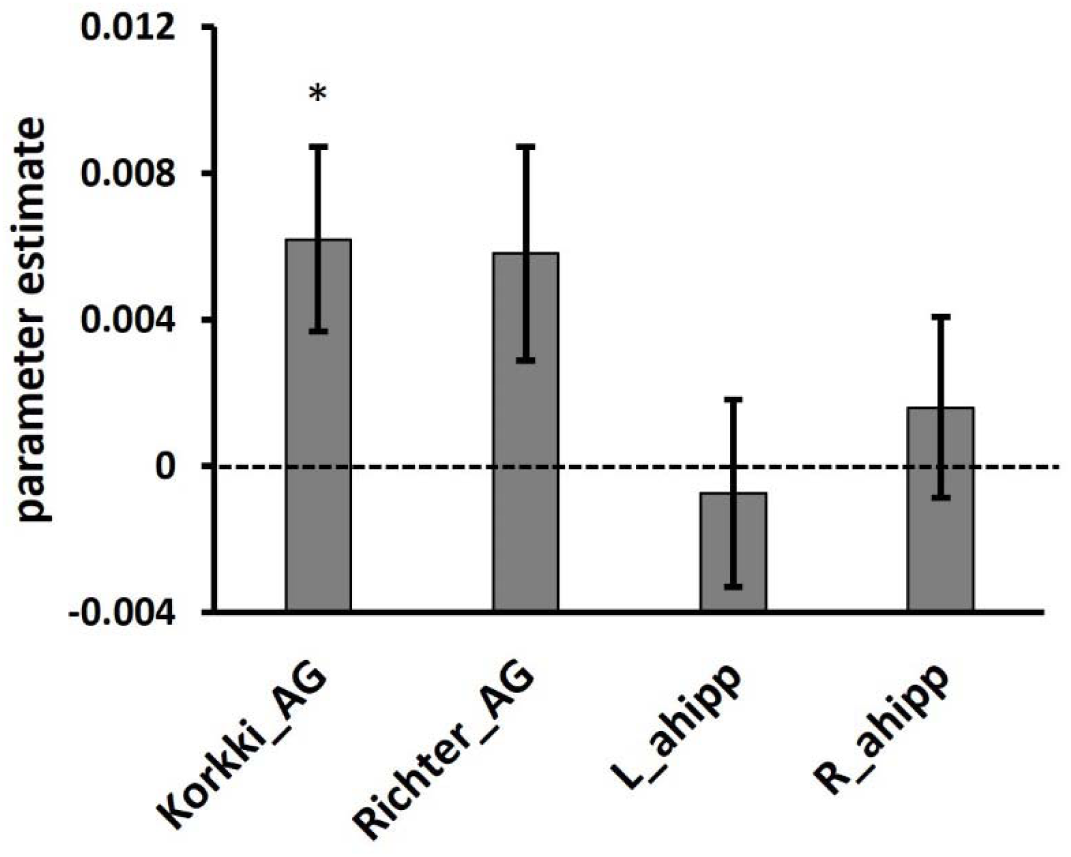
Parameter estimates for the parametric regressors tracking memory precision derived from the angular gyrus and hippocampal ROIs. Error bars indicate standard errors of the mean. Korkki_AG: angular gyrus ROI based on the peak reported by Korkki et al. (2023); Richter_AG: angular gyrus ROI based on the peak reported by Richter et al. (2016); ahipp: anterior hippocampus. *p < 0.05.

Turning to retrieval success effects, no significant differences were identified between location hit and guess trials in any ROI (all ps > 0.46, see Figure 6). A significant ‘novelty’ effect, as reflected by greater activity for CRs than for guess or hit trials, was however evident in each case (ps < 0.033, Cohen’s ds > 0.47).

**Figure 6.**
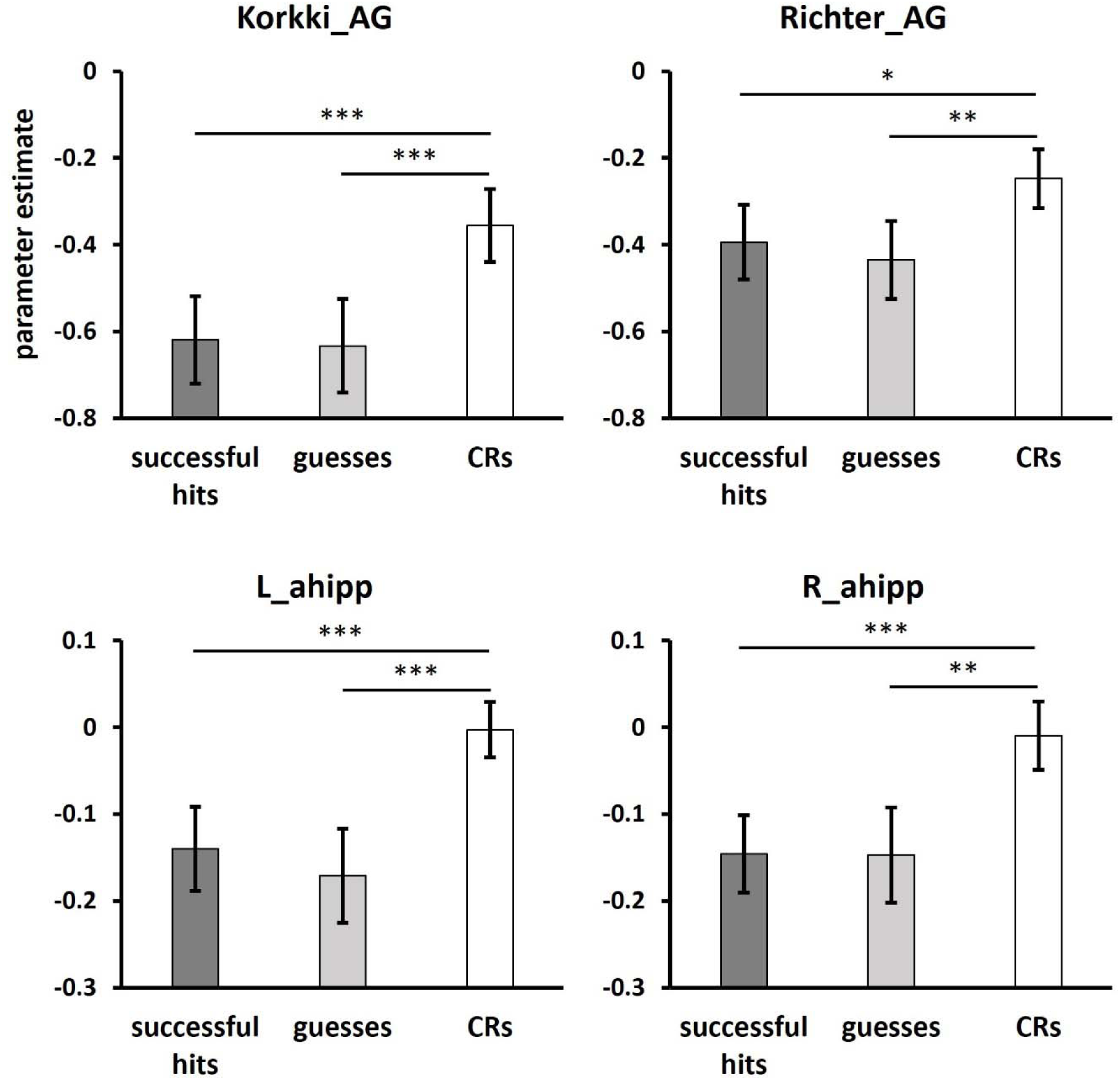
Parameter estimates extracted from the angular gyrus and the hippocampal ROIs. Error bars indicate standard errors of the mean. Korkki_AG: angular gyrus ROI based on the peak reported by Korkki et al. (2023); Richter_AG: angular gyrus ROI based on the peak reported by Richter et al. (2016); ahipp: anterior hippocampus. *p < 0.05; **p < 0.01; ***p < 0.001.

Findings from the contrast between high-precision and low-precision hit trials were highly consistent with the results reported above (Figure 7). Specifically, the contrast between high-precision and low-precision hit trials was significant in both the Korkki (t_22_ = 2.38, p = 0.026, Cohen’s d = 0.50) and Richter ROI (t_22_ = 2.10, p = 0.047, Cohen’s d = 0.44). There was no significant difference between the high vs. low precision hit trials in the hippocampus (ps > 0.44). In the case of retrieval success, no ROI demonstrated a significant effect (ps > 0.45). The findings remained nonsignificant when the contrast was between high-precision hits and guesses (ps > 0.16).

**Figure 7.**
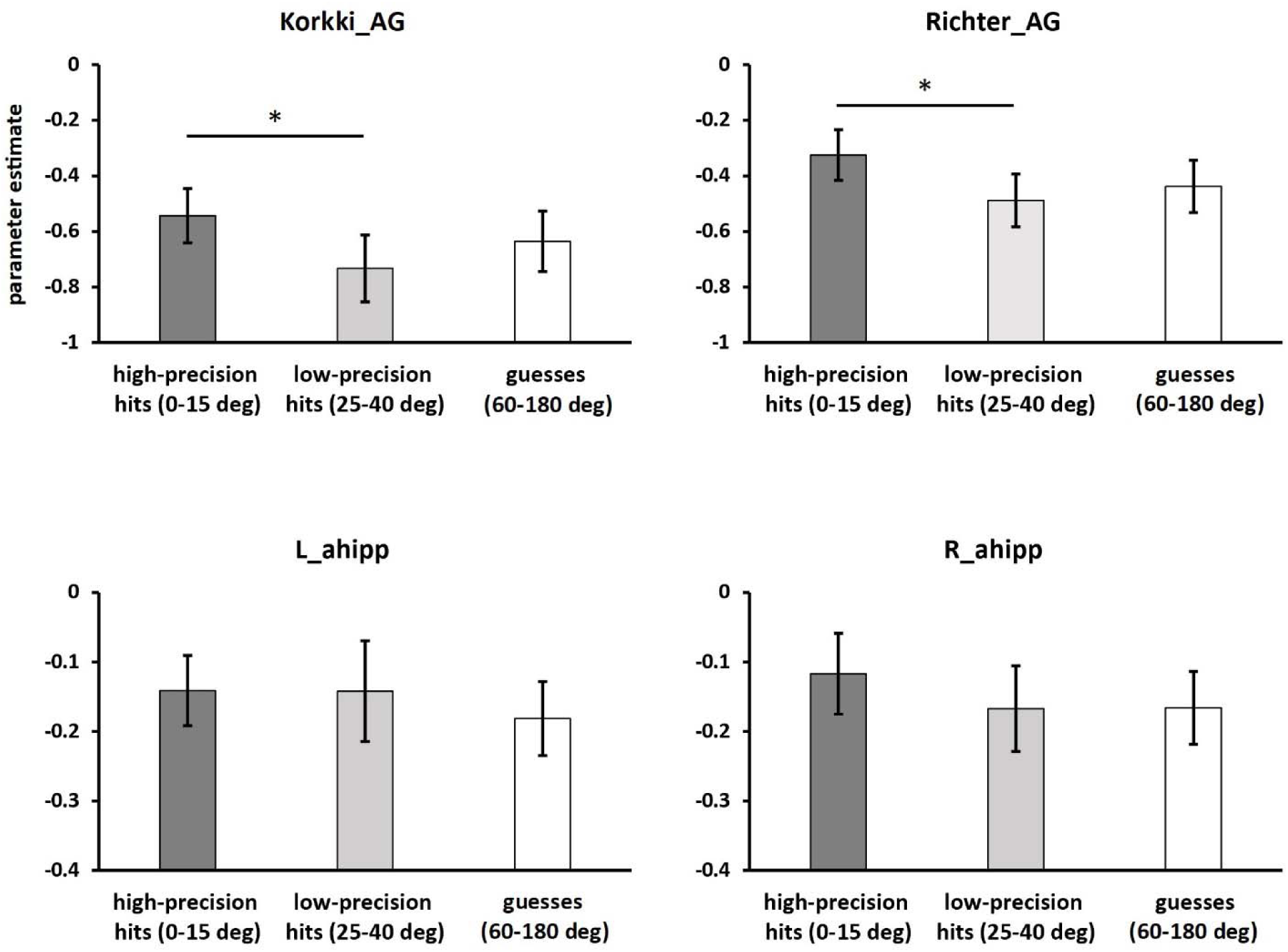
Parameter estimates of high-, low-precision hits and guess trials extracted from the angular gyrus and the hippocampal ROIs. Error bars indicate standard errors of the mean. Korkki_AG: angular gyrus ROI based on the peak reported by Korkki et al. (2023); Richter_AG: angular gyrus ROI based on the peak reported by Richter et al. (2016); ahipp: anterior hippocampus. *p < 0.05.

#### 3.2.3 PSA

A reliable item-level reinstatement effect was identified in the AG ROI for the high-precision trials (t_22_ = 4.36, p < 0.001, Cohen’s d = 0.91). No effect was evident however for either the low-precision or guess trials (ps > 0.12, see Figure 8). Additionally, reinstatement was significantly greater for the high-precision hits than for either low-precision hits (t_22_ = 2.22, p = 0.037, Cohen’s d = 0.46) or guesses (t_22_ = 2.18, p = 0.040, Cohen’s d = 0.45). There was no evidence of item-level reinstatement in the left or right anterior hippocampus for any class of memory judgment (ps > 0.086).

**Figure 8.**
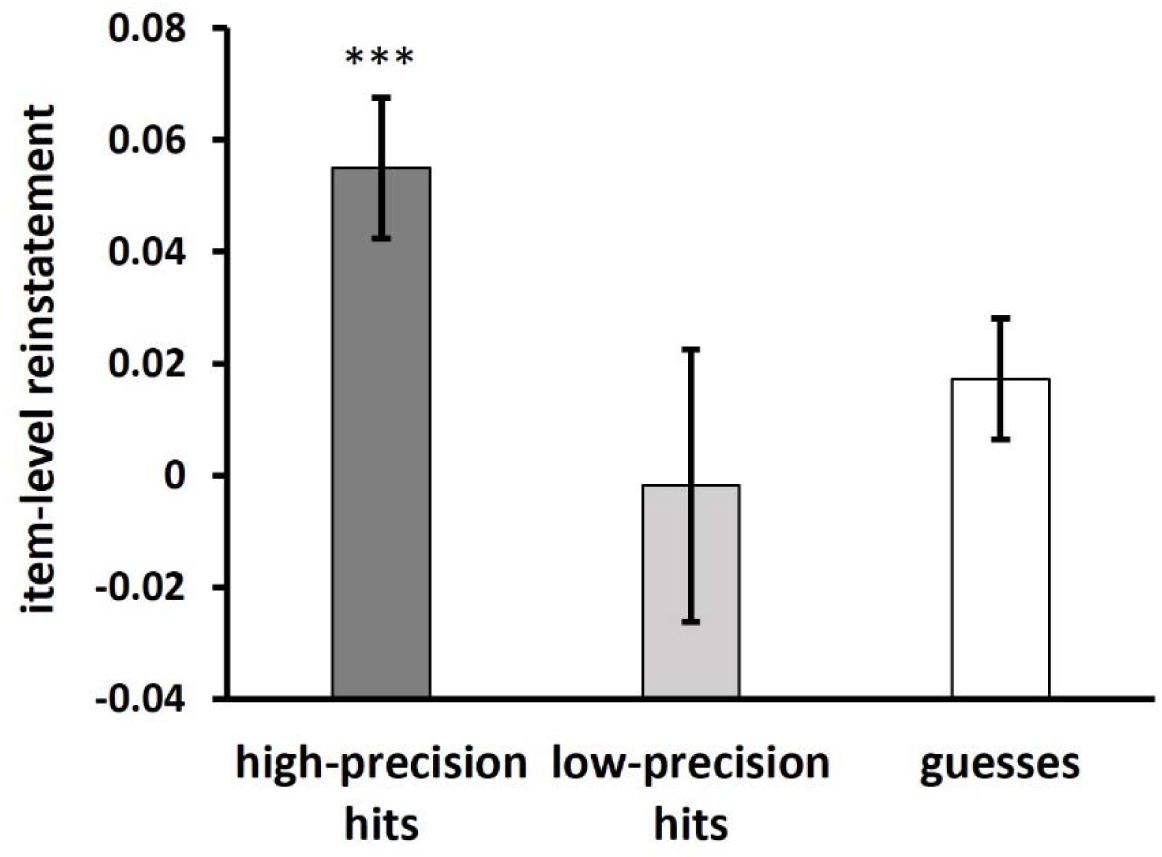
Estimates of item-level reinstatement in the anatomically defined AG. ***p < 0.001.

## 4. Discussion

We examined neural correlates of retrieval success and memory precision in a sample of healthy young adults. At the whole brain level we were unable to identify any region at the pre-experimental threshold where BOLD activity covaried with precision. However, the more sensitive ROI approach identified an AG region where activity did demonstrate a precision effect. Complementing these findings, using PSA we identified reliable item-level reinstatement effects in the AG for high-precision memory judgments, whereas no such effects were evident for low-precision or guess trials. Regardless of the analysis approach, hippocampal activity was insensitive to both retrieval success and precision. Additionally, a whole-brain analysis identified several regions where activity elicited by test items differed according to their study history. We expand on these findings below.

In the present study, we used a positional response accuracy procedure similar to that employed in early studies of memory precision (e.g. Harlow and Donaldson, 2014). The experimental design differs from prior fMRI studies examining precision and retrieval success (Cooper et al., 2017; Korkki et al., 2023; Richter et al., 2016) in two main ways. First, in the prior experiments, study items were superimposed on complex scene backgrounds, and precision was operationalized by accuracy of retrieval of multiple features of the study item [location, orientation and color in Richter et al. (2016) and Cooper et al. (2017); location and color in Korkki et al. (2023)]. In the present case, however, study items were presented against a uniform background and memory for a single contextual feature (location) was probed. Second, unlike in prior fMRI and behavioral studies, the test phase included both studied and unstudied items, and participants were required to discriminate between these two classes of test item. Thus, whereas in prior studies precision estimates were derived from an unknown mixture of recognized and unrecognized items, in the present case estimates were restricted to items that participants endorsed as having been studied. As we discuss below, these two aspects of our experimental procedure may have contributed to our failure to identify retrieval success effects in the AG and hippocampus.

Motivated by the inconsistent prior evidence pertaining to the sensitivity of the AG and hippocampus to memory precision and retrieval success (Cooper et al., 2017; Korkki et al., 2023; Richter et al., 2016), we examined precision- and retrieval-related activity in these regions. Consistent with the prior findings (Cooper et al., 2017; Korkki et al., 2023; Richter et al., 2016), the results suggest that BOLD activity in a small region of the left AG is positively correlated with a trial-wise measure of memory precision, while providing no evidence of a retrieval success effect in this region. This pattern of findings was buttressed by the results of a categorical analysis that also indicated that fMRI responses in the left AG were greater for high-than for low-precision location hits, but did not demonstrate a difference between location hits and guesses. In a complementary analysis, PSA revealed that the AG demonstrated a robust item-specific reinstatement effect that was exclusive to high-precision memory judgments. These findings are consistent with the idea that the judgments were supported by the encoding and subsequent retrieval of item-specific information that was diagnostic of the studied locations. Together, the present univariate and PSA findings align with prior reports from other studies employing univariate (e.g. Mayes et al., 2019; Thakral et al. 2015; Vilberg and Rugg, 2009a,b; Yu et al., 2012a) and multi-voxel analysis approaches (e.g. Kuhl and Chun, 2014; Lee et al., 2019) indicating that fMRI activity in the left AG is sensitive to the strength and fidelity of the memory signal supporting recollection-related memory judgments (see Rugg and King, 2018 for review).

That being said, we could find no evidence that univariate AG activity differed between high precision trials (or location hits more generally) and guesses; that is, there was no evidence of a univariate ‘recollection effect’ in this region (see Figure 7). On its face, this finding seems to be inconsistent with a role for the AG in signaling memory precision – by definition, guess trials are trials on which memory precision is nominally zero. A possible resolution of this seeming paradox is to assume that guess trials included trials on which recollection about the study event was successful, and hence was associated with enhanced AG activity, but where the recollected information was non-diagnostic of location (cf. Yonelinas and Jacoby, 1996). This account however raises an additional problem, given that the retrieval of non-diagnostic information arguably would be equally likely for high-precision hit trials. If so, it remains unclear why high-precision location hits failed to elicit greater AG activity than guesses. A possible solution to this conundrum is to assume that whereas location hit trials reflected study processing that focused on the encoding of item-location associations, guess trials arose when participants attended to some other aspect of the study event at the expense of location information (as noted above, in contrast to prior fMRI studies, here guess trials were restricted to test items that carried a memory signal sufficient to support an accurate recognition judgment). In other words, there was a trade off between the encoding of diagnostic and non-diagnostic information. Future research will be required to examine the validity of this admittedly speculative and post-hoc account.

In contrast to the findings for the left AG, we were unable to identify either retrieval success or precision effects in the hippocampus. Nor were we able to find evidence from PSA of reinstatement effects analogous to those identified in the AG (see above), a null finding that held both for the whole hippocampus as well as its anterior extent. Our results thus stand in contrast to prior reports that hippocampal activity and multi-voxel reinstatement are enhanced when recollection is successful (e.g. de Chastelaine et al., 2016; Dulas and Duarte, 2016; Tompary et al., 2016) as well as reports of hippocampal retrieval success effects in prior fMRI studies of memory precision (Richter et al., 2016; Cooper et al., 2017; Korrki et al., 2023). We consider three possible explanations for these null findings, which are not necessarily mutually exclusive.

One possible explanation is essentially the same as that advanced to explain the absence of AG retrieval success effects: that is, guess trials were associated with the recollection of non-diagnostic information that attenuated any difference in the BOLD activity elicited by hits vs. guesses. A second possible explanation is that location hit trials were supported by a relatively weak recollection signal. Prior findings indicate that, like activity in the AG, hippocampal activity covaries with the amount of retrieved contextual information, and that recollection effects in this region are undetectable when only a small amount of information is recollected (Mayes et al., 2019; Thakral et al., 2015; Yu et al., 2012b; for review, see Rugg et al., 2012). From this perspective, it might be relevant that, as already noted, prior studies that reported hippocampal retrieval success and precision effects employed complex, content-rich study events (object-scene pairings) that likely led to the retrieval of similarly content-rich memories. A final possibility is that the voxel size used in the present experiment (2.5 mm^3^) was too large to allow the identification of effects that might be restricted to a single hippocampal subfield [we note however that the prior studies in which univariate hippocampal retrieval success and precision effects were reported (Cooper et al., 2017; Korkki et al., 2023; Richter et al., 2016) employed a larger voxel size than that used in the present study, and that multi-voxel reinstatement has been reported previously at the level of the whole hippocampus (Tompary et al., 2016)]. These issues can be addressed in future studies with higher resolution fMRI sequences that target the hippocampus.

In both the AG and the hippocampus BOLD activity elicited by CRs was robustly greater than that elicited either by location hits or guess trials. These findings are difficult to interpret. They might reflect processes related to the detection of novel information and the subsequent initiation of memory encoding operations (see, for example, Lisman and Grace, 2005; Nyberg, 2005). Alternatively, they might reflect a task difficulty or time on task effect, given that participants likely expended less effort during the processing of unstudied than studied test items (cf. Humphreys and Lambon Ralph, 2015).

Turning to the whole brain analyses, we identified robust effects of trial type in a variety of cortical regions. However, in only three of these regions – left and right DPC and right premotor cortex - was there evidence of a memory retrieval effect, such that activity varied in a graded manner between location hits, guesses and CRs (Figure 4). Intriguingly, left DPC has consistently been implicated in familiarity-driven recognition memory (see Rugg and King, 2018, for review). From this perspective, the present finding might indicate a role for familiarity in both item memory and successful location judgments (see Yonelinas, 1999 for evidence that familiarity can support source memory) or, alternately, that the encoding conditions that are conducive to successful location memory also act to strengthen item familiarity.

In addition, both left and right DPC, together with premotor cortex and the DLPFC, demonstrated greater activity for location hits and guesses than for correct rejection trials. Rather than reflecting mnemonic processing, these findings might be attributed to the differential visuomotor demands associated with the different memory judgments. Notably, each of these regions has been implicated in visuomotor function. For instance, the DPC has been implicated in the transformation of information from a sensory reference frame into a motor reference frame in support of visually guided action (Cohen and Anderson, 2002; Culham and Valyear, 2006). It seems likely that these disparate regions interacted to support the visuomotor demands of the positional response accuracy task, which required fine-grained, visually-guided motor control. From this perspective, the greater activity elicited on location hit and guess trials than on CR trials is a reflection of the greater visuomotor demands associated with memory-guided rather than randomly determined placement of the test cursor.

The present study has three main limitations. First, the sample size was relatively small, signaling the need for caution in accepting null findings, especially in respect of the hippocampal retrieval success and precision effects. Second, we only assessed location memory, and hence we were unable to estimate the influence of the retrieval of non-diagnostic information on the BOLD activity elicited by studied test items. Future studies should acquire trial-wise subjective reports about recollected content in addition to precision estimates. Finally, the validity of the mixture model employed here in the analysis of the behavioral data has been questioned on the grounds that distributions of continuous error metrics can be fit equally well or better by a single-component model (Schurgin et al., 2020). As was noted by Berens et al. (2020), however, this seemingly more parsimonious model has largely been applied to studies of short-term and working memory, and cannot easily accommodate the double dissociations between success and precision that have been reported in the long-term memory literature (see Introduction).

In conclusion, in line with prior reports, we found that neural activity in the angular gyrus was sensitive to memory precision but not to retrieval success. The findings are consistent with other evidence suggesting that retrieval success and precision are dissociable features of memory retrieval.

## Acknowledgments

This work was supported by the National Institute of Neurological Disorders and Stroke (grant number R01NS114913). We thank our experimental participants for their participation.

